# IID-KG: An ontology-aligned literature-derived knowledge graph for infectious and immune-mediated diseases

**DOI:** 10.64898/2026.05.21.727015

**Authors:** Feng Pan, Yuan Zhang, Jianan Wang, Ming-Chieh Liu, Xin Sui, Han Yue, Jinfeng Zhang

## Abstract

Infectious and immune-mediated diseases (IIDs) represent a broad and rapidly expanding biomedical literature domain in which scalable evidence extraction, disease ontology refinement, and interpretable knowledge integration are essential for biomedical discovery. We constructed an IID-specific biomedical knowledge graph (IID KG) from PubMed abstracts and PMC full-text articles by integrating nested named entity recognition, ontology-guided identifier assignment, full-text relation extraction, and relation-resolution strategies. A gold-standard corpus of 500 PubMed abstracts and 8 PMC full-text articles was manually annotated for nested biomedical entities across six entity types. The resulting models were applied to 30,128,068 PubMed abstracts and 1,385,500 IID-related PMC full-text articles. A unified IID ontology was developed from 411,341 disease terms using hierarchical text classification, large language model-based refinement, ontology cross-referencing, and expert review, yielding 179,657 confirmed MeSH mappings. The final IID KG contains approximately 1,837,513 unique entities and 16,295,390 unique relations across eight relation types. The resource was released publicly together with repurposing workflows, supporting ontology-aligned literature mining, disease mechanism analysis, and drug-repurposing hypothesis generation for IID research.

## 1. Introduction

The biomedical literature has become one of the largest and most rapidly expanding sources of mechanistic, translational, and clinical evidence. For infectious and immune-mediated diseases (IIDs), this growth creates a particularly acute information-integration problem: relevant observations are distributed across millions of PubMed abstracts and full-text articles, use heterogeneous disease names and synonyms, and often describe relationships among diseases, genes, chemicals, variants, species, and cell lines in unstructured language. Manual review cannot scale to this volume, and keyword-based retrieval alone is insufficient when the goal is to identify structured evidence, connect related findings across studies, and generate testable hypotheses for disease mechanisms, drug repurposing, or target discovery [1].

Knowledge graphs (KGs) offer a practical framework for converting unstructured biomedical text into computable evidence. In a biomedical KG, normalized entities are represented as nodes and their relationships are represented as typed edges, allowing direct evidence retrieval, cross-article aggregation, and downstream reasoning over multi-hop paths. Recent large-scale biomedical KG work has shown that literature-derived information extraction pipelines can be applied across PubMed-scale corpora and integrated with structured biomedical databases to support downstream AI and discovery tasks [2]. Such graphs are especially useful for IID research because infectious and immune-mediated conditions frequently involve complex interactions among host genes, pathogens, immune pathways, drugs, genetic variants, and disease phenotypes. However, building a high-quality literature-derived KG for this domain requires more than large-scale text mining. It requires reliable named entity recognition (NER), robust entity normalization, accurate relation extraction, explicit treatment of novelty, and an ontology that can resolve disease synonyms and distinguish true disease entities from symptoms, organisms, vaccines, broad descriptors, or other non-disease mentions [3].

Recent community challenges and biomedical NLP benchmarks have accelerated the development of methods for NER, relation extraction, entity normalization, and end-to-end KG construction. The LitCoin NLP Challenge and BioCreative BioRED tasks established annotation schemas that include common biomedical entity types and relation categories relevant to translational research, including associations, binding, conversion, cotreatment, drug interactions, positive correlations, and negative correlations [4]. The central deliverable of the present work is a dedicated IID-specific KG rather than a generic enhancement of an existing graph. The IID KG was designed to focus the literature-mining pipeline, disease ontology refinement, and downstream reasoning around infectious and immune-mediated disease research.

A major barrier to constructing such a graph is disease ontology quality. Disease mentions in biomedical literature are highly variable: the same concept may appear under different synonyms, abbreviations, or spelling variants, whereas related but distinct concepts may be incorrectly grouped together. Biomedical entity normalization aims to map textual mentions to standard identifiers in resources such as MeSH or UMLS, and prior work emphasizes that synonymy, abbreviation, ambiguity, and incomplete dictionaries make this step difficult [5]. There are cases such as disease names being confused with pathogens, broad genus-level terms, combination vaccines, or related infections. These errors can propagate directly into KG construction because relation extraction operates over normalized entity pairs. If synonyms are split incorrectly, evidence is fragmented; if distinct concepts are merged incorrectly, spurious relations are introduced. Therefore, IID KG construction requires a disease-focused ontology that combines MeSH alignment, external ontology cross-referencing, machine-learning classification, LLM-assisted refinement, and expert manual validation.

Another challenge is the use of full-text articles. Abstracts are compact and information-dense, but many mechanistic and experimental details are found only in full texts. Full-text mining can improve coverage, yet it introduces substantial heterogeneity across article sections. Methods sections often contain procedural language, Results sections contain dense quantitative and experimental descriptions, and Discussion or Conclusion sections may summarize broader biological implications. A KG construction pipeline must therefore evaluate NER and relation extraction not only at the document level but also across full-text sections, and must resolve multiple or conflicting sentence-level predictions for the same entity pair within long articles [6].

In this study, we constructed and released a standalone IID-specific biomedical knowledge graph from PubMed abstracts and IID-related PMC full-text articles. The workflow included: (i) creation of a gold-standard nested entity annotation corpus from PubMed abstracts and PMC full texts; (ii) development and evaluation of abstract-level and full-text nested NER models; (iii) large-scale processing of more than 30 million PubMed abstracts and nearly 1.4 million IID-related PMC full texts; (iv) construction and refinement of a unified IID ontology using hierarchical classification, LLM-based mapping, ontology cross-referencing, and manual review; (v) ontology-aligned relation extraction and full-text relation resolution; and (vi) integration of the resulting normalized entities and relation triples into an open IID KG suitable for evidence retrieval, hypothesis generation, and PSR-based drug repurposing workflows.

## 2. Materials and Methods

### 2.1. Data acquisition and corpus preparation

All available PubMed abstracts up to April 14, 2025 were retrieved, yielding 30,128,068 records. These abstracts were processed using the finalized PubMed NER pipeline and filtered for infectious and immune-mediated disease mentions using the unified IID ontology. This filtering identified 6,328,681 IID-related PubMed abstracts. PubMed-PMC linkage information was then used to retrieve corresponding PMC full-text articles, resulting in 1,385,500 IID-related PMC full texts. Full texts were initially obtained in XML format and converted to JSON to support downstream NER, identifier assignment, relation extraction, and large-scale integration.

### 2.2. Gold-standard nested entity annotation

A gold-standard nested entity corpus was created to support model training, cross-domain evaluation, and full-text adaptation. The corpus included 500 PubMed abstracts and 8 PMC full-text articles manually annotated for six biomedical entity types: Disease, Gene/GeneProduct, Chemical, DNAmutation, CellLine, and Species. Annotation was performed by trained biomedical annotators using Insilicom’s Text Annotation and Modeling (ITAM) platform, which supports text navigation, span selection, schema-driven entity type assignment, and visualization of nested or overlapping spans. All documents were double annotated, disagreements were adjudicated by a senior annotator, and inter-annotator agreement was periodically evaluated to monitor consistency.

### 2.3. Nested NER model development and evaluation

Two NER configurations were evaluated. The abstract-level model was trained exclusively on the 500 annotated PubMed abstracts to assess in-domain performance and cross-domain transfer to PMC full texts. The full-text model was trained on a combined dataset consisting of 500 annotated PubMed abstracts and annotated PMC full texts, with 6 full texts used for training and 2 full texts held out for testing in the reported configuration. Performance was measured using true positives, false positives, false negatives, precision, recall, and F1 score. Section-wise evaluation was conducted for PMC articles to quantify performance differences across Introduction, Methods, Results, and Conclusion sections. External robustness was evaluated on the Europe PMC annotated corpus for Disease and Gene entity types; for Disease, Condition mentions were included in gold annotations to align with the internal schema.

Following inference, document-level post-processing was applied to improve entity type consistency within each article. Repeated mentions of the same entity were identified, type conflicts were resolved by assigning the most frequently occurring type, and the BIO tagging scheme was preserved. The post-processed entity outputs were used for relation extraction and knowledge graph integration.

### 2.4. Relation extraction and relation resolution

Relation extraction was performed on sentence pairs co-mentioning the same entity pair. Sentence-level predictions were aggregated for each entity pair, and retained non-NOT relations were prepared for novelty prediction by concatenating the top-ranked supporting sentences within the BERT token limit. A novelty-based filtering strategy was then applied: when at least one novel relation existed for an entity pair, only novel relations were retained; otherwise, all non-novel relations were retained. Remaining candidate relations were ranked by informativeness using the priority hierarchy Positive correlation and Negative correlation, followed by Bind and Conversion, followed by Cotreatment, Comparison, and Drug Interaction, followed by Association. Ties within the top-priority group were resolved by the highest combined relation probability and then by the highest novelty probability when needed. An optional abstract-level override prioritized conflicting novel abstract-level evidence for the same entity pair.

### 2.5. IID ontology development and disease identifier assignment

A unified IID ontology was developed to normalize disease mentions and support consistent integration into the newly constructed IID KG. IID-related disease terms were extracted from IKraph and combined with candidates from PubMed and PMC NER outputs. Candidate MeSH identifiers were assigned using a Hierarchical Text Classification (HTC) model HiDEC trained on MeSH tree nodes [7]. For terms not directly present in MeSH, additional candidate identifiers were generated from model predictions, existing IKraph assignments, and LLM-based questioning, followed by Claude 3.7-based selection of the most appropriate mapping.

Noise filtering was performed to remove non-disease terms such as symptoms, species names, and general conditions by cross-referencing MeSH, MONDO, and Disease Ontology resources and by applying GPT-4o-based classification. Disease terms without MeSH identifiers were harmonized using all-MiniLM-L6-v2 semantic similarity [8], Claude 3.7 synonymy validation, and a custom ontology ID merging algorithm. Synonym mappings within MeSH descriptors were refined by identifying disease terms likely to map to more specific MeSH concept IDs rather than broader descriptor IDs; approximately 6,000 candidate terms were manually inspected after LLM-based reassignment suggestions. Assigned identifiers were categorized into five validation levels: manually annotated terms, direct MeSH mappings, LLM-confirmed IDs, post-processed refinements, and newly created IDs for unmapped terms.

### 2.6. Knowledge graph construction and downstream demonstration

Ontology-aligned entities and relation triples extracted from PubMed abstracts and PMC full texts were integrated into a standalone IID-specific knowledge graph. Nodes represent normalized biomedical entities, including diseases from the unified IID ontology, genes and gene products, chemicals, DNA mutations, cell lines, and species. Edges represent ontology-aligned biomedical relations extracted from abstracts and full texts, including mechanistic, correlation-based, treatment-related, and general association relations. The resulting IID KG was designed as a dedicated resource for infectious and immune-mediated disease research while remaining interoperable with IKraph-derived workflows.

## 3. Results

### 3.1. Gold-standard corpus and abstract-level NER performance

The abstract-level model showed strong in-domain performance on PubMed abstracts, with an F1 score of 0.8669. When applied to PMC full-text articles, performance decreased to 0.8223, consistent with the greater length, structural complexity, and linguistic variability of full-text biomedical articles.

Section-wise evaluation on the eight PMC full-text articles showed the highest performance in Conclusion and Results sections and the lowest performance in Methods. This pattern indicates that full-text NER performance is section dependent and that structurally complex methodological language remains a key source of extraction difficulty.

### 3.2. Full-text model performance and external validation

The full-text model trained on the combined abstract and full-text corpus improved robustness for full-text processing. In the held-out PMC evaluation configuration, it achieved an overall F1 score of 0.8418, with particularly improved performance in the Methods section relative to the abstract-only model.

External evaluation on the Europe PMC annotated corpus confirmed robust recognition of Disease and Gene entities across biomedical full-text sections. Disease recognition achieved an overall F1 score of 0.9305, with precision and recall above 0.92 overall. Gene recognition achieved an overall F1 score of 0.8437, with stronger performance in Abstract, Introduction, Discussion, and Conclusion sections and lower performance in Methods, where gene-rich experimental descriptions introduce greater lexical diversity.

### 3.3. Large-scale processing outcomes

The finalized pipelines were deployed at large scale. All 30,128,068 PubMed abstracts up to April 14, 2025 were processed with the abstract-level NER model, and 6,328,681 abstracts containing IID mentions were identified through ontology-based filtering. Corresponding PMC full-text retrieval yielded 1,385,500 IID-related full-text articles. These full texts were converted from XML to JSON, processed with the full-text NER model, post-processed for document-level entity consistency, and submitted to relation extraction and relation resolution. Parallelized distributed processing, logging, and error-handling mechanisms enabled completion of the large-scale abstract and full-text processing within the project period.

### 3.4. IID ontology refinement and validation

Hierarchical text classification supported initial candidate MeSH assignment for infectious and immune-related disease terms. The infectious disease classifier achieved strong test F1 performance across tested encoders, with the best reported F1 of 0.9633 using bert-base-uncased. The immune-related disease classifier achieved a test F1 of 0.956 using bert-base-uncased (Table 6-7).

**Table 1.** NER model performance on PubMed abstracts and PMC full-text articles.

| Dataset | TP (N_entity) | FP (N_entity) | FN (N_entity) | Precision | Recall | F1 |
| --- | --- | --- | --- | --- | --- | --- |
| PubMed Abstract | 4040 | 598 | 638 | 0.8720 | 0.8625 | 0.8669 |
| PMC (8 total) | 5527 | 1257 | 1132 | 0.8147 | 0.8300 | 0.8223 |

**Table 2.** Section-wise NER performance on 8 PMC full-text articles.

| Section | TP | FP | FN | Precision | Recall | F1 |
| --- | --- | --- | --- | --- | --- | --- |
| Introduction | 818 | 170 | 184 | 0.8279 | 0.8164 | 0.8221 |
| Methods | 351 | 141 | 92 | 0.7134 | 0.7923 | 0.7508 |
| Results | 3713 | 831 | 761 | 0.8171 | 0.8299 | 0.8235 |
| Conclusion | 645 | 115 | 95 | 0.8487 | 0.8716 | 0.86 |

**Table 3.** NER performance of the full-text model on PMC full-text articles.

| Precision | Recall | F1 | F1 (Introduction) | F1 (Methods) | F1 (Results) | F1 (Conclusion) |
| --- | --- | --- | --- | --- | --- | --- |
| 0.8569 | 0.8272 | 0.8418 | 0.8605 | 0.7789 | 0.8387 | 0.8739 |

**Table 4.** Disease entity recognition performance on the Europe PMC annotated corpus.

| Section | TP | FP | FN | Precision | Recall | F1 |
| --- | --- | --- | --- | --- | --- | --- |
| Abstract | 1374 | 81 | 73 | 0.9443 | 0.9496 | 0.9469 |
| Intro | 3302 | 165 | 140 | 0.9524 | 0.9593 | 0.9559 |
| Methods | 1997 | 232 | 118 | 0.8959 | 0.9442 | 0.9194 |
| Results | 4071 | 401 | 455 | 0.9103 | 0.8995 | 0.9049 |
| Discuss | 4542 | 362 | 293 | 0.9262 | 0.9394 | 0.9327 |
| Conclusion | 362 | 11 | 8 | 0.9705 | 0.9784 | 0.9744 |
| All | 15648 | 1252 | 1087 | 0.9259 | 0.9350 | 0.9305 |

**Table 5.** Gene entity recognition performance on the Europe PMC annotated corpus.

| Section | TP | FP | FN | Precision | Recall | F1 |
| --- | --- | --- | --- | --- | --- | --- |
| Abstract | 1602 | 343 | 110 | 0.8237 | 0.9357 | 0.8761 |
| Intro | 3010 | 627 | 225 | 0.8276 | 0.9304 | 0.8760 |
| Methods | 3603 | 1221 | 701 | 0.7469 | 0.8371 | 0.7894 |
| Results | 9645 | 2842 | 975 | 0.7724 | 0.9082 | 0.8348 |
| Discuss | 6313 | 1534 | 424 | 0.8045 | 0.9371 | 0.8657 |
| Conclusion | 396 | 79 | 24 | 0.8337 | 0.9429 | 0.8849 |
| All | 24569 | 6646 | 2459 | 0.7871 | 0.9090 | 0.8437 |

**Table 6.** Dataset statistics and performance metrics for infectious disease classification.

| N of nodes | max depth | train | dev | test | pretrained encoder | test F1 |
| --- | --- | --- | --- | --- | --- | --- |
| 1410 | 9 | 4922 | 615 | 615 | bert-base-uncased | 0.9633 |
| 1410 | 9 | 4922 | 615 | 615 | pubmed-bert-base-uncased-abstract | 0.9162 |
| 1410 | 9 | 4922 | 615 | 615 | bert-base-cased-finetuned-ner-NCBI_Disease | 0.921 |

**Table 7.** Dataset statistics and performance metrics for immune-related disease classification.

| N of nodes | max depth | train | dev | test | pretrained encoder | test F1 |
| --- | --- | --- | --- | --- | --- | --- |
| 279 | 9 | 3116 | 390 | 390 | bert-base-uncased | 0.956 |

Across the ontology pipeline, 411,341 disease terms were processed, yielding 179,657 terms with confirmed MeSH ID mappings after LLM refinement (Fig. 3). Noise filtering achieved F1 scores of 0.9351 for infectious diseases and 0.9639 for immune-related diseases. Manual review covered 3,237 infectious and 2,462 immune-related terms, including all terms with at least 1,000 PubMed mentions. Comparative evaluation against MeSH, MONDO, and Disease Ontology showed strong MeSH alignment and broad infectious disease coverage, while also revealing category definition differences for immune-related diseases across external resources (Table 8-9).

**Table 8.** Comparison of ID and synonym counts for infectious diseases between the unified ontology and external databases.

| Source | Infectious IDs | IDs Found in IKraph | IDs Classified as Infectious in IKraph | Infectious Synonyms | Synonyms Found in IKraph | Synonyms Classified as Infectious in IKraph |
| --- | --- | --- | --- | --- | --- | --- |
| MeSH | 783 | 746 | 746 | 6,935 | 2,887 | 2,887 |
| MONDO | 1,074 | 1,023 | 942 | 3,544 | 1,964 | 1,682 |
| DO | 528 | 498 | 484 | 1,401 | 970 | 898 |

**Table 9.**
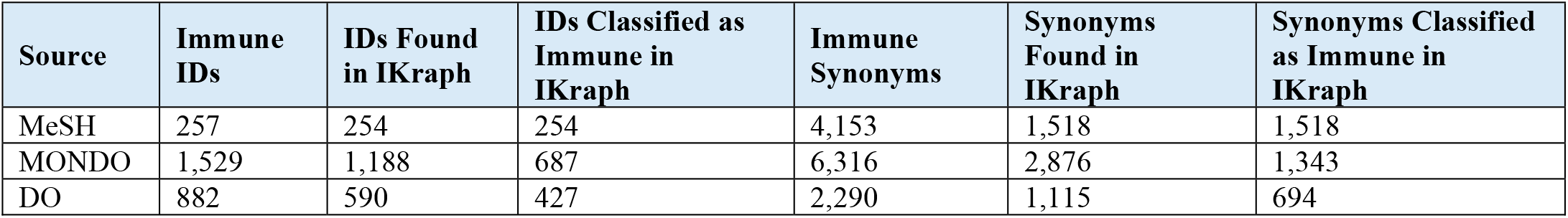
Comparison of ID and synonym counts for immune-related diseases between the unified ontology and external databases.

**Table 10.** Distribution of relation instances in the IID-specific knowledge graph by entity pair type.

|  | C-D | V-G | V-V | C-V | C-C | D-G | G-G | V-D | C-G |
| --- | --- | --- | --- | --- | --- | --- | --- | --- | --- |
| Association | 91442 | 65389 | 1193 | 22264 | 385033 | 524114 | 3029149 | 7044 | 3407503 |
| Bind | 61 | 457 | 11 | 154 | 96457 | 313 | 396656 | 3 | 755312 |
| Comparison | 96 | 4 | 13 | 81 | 218314 | 3 | 1585 | 0 | 7401 |
| Conversion | 0 | 0 | 0 | 0 | 13835 | 0 | 9 | 0 | 433 |
| Cotreatment | 325 | 15 | 22 | 81 | 453918 | 90 | 86837 | 2 | 19983 |
| Drug Interaction | 0 | 0 | 0 | 0 | 12581 | 0 | 3008 | 0 | 8660 |
| NegativeCorrelation | 211582 | 8958 | 266 | 4267 | 594214 | 67015 | 2161444 | 340 | 947642 |
| PositiveCorrelation | 38389 | 8085 | 119 | 2948 | 235451 | 61273 | 1300065 | 474 | 1042677 |

**Figure 1.**
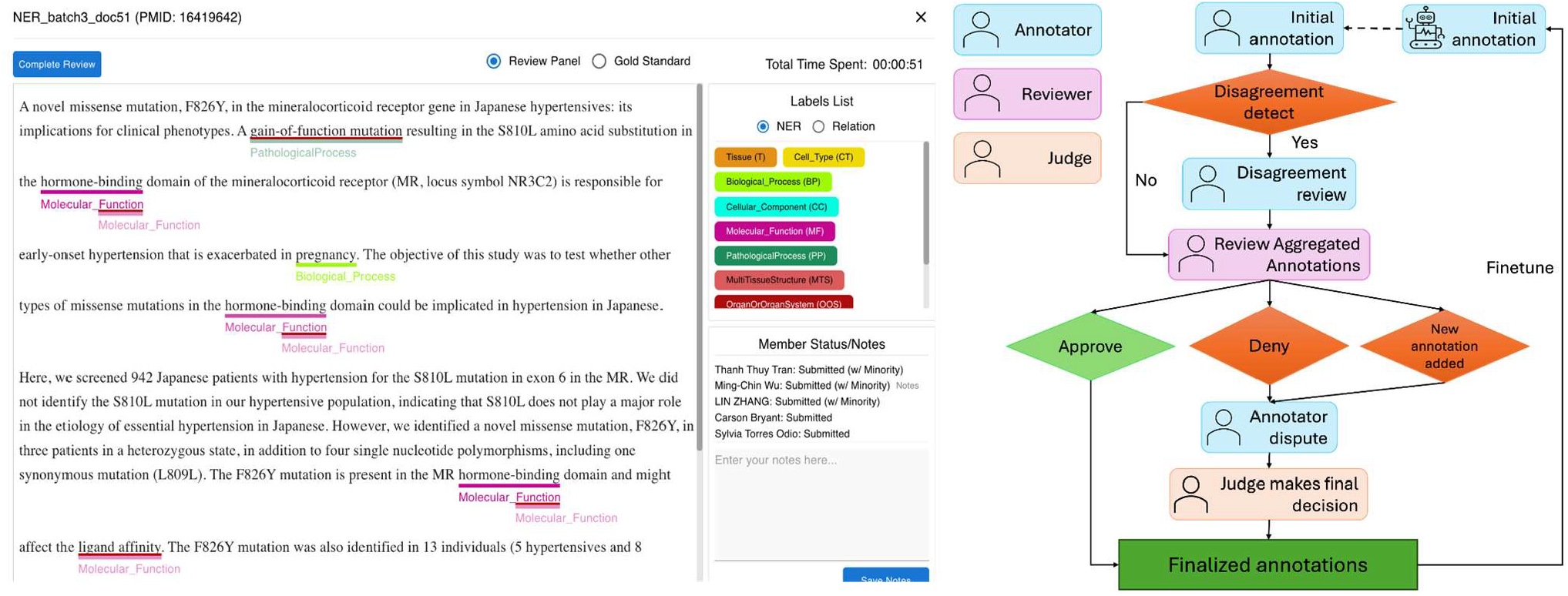
ITAM annotation platform and annotation workflow. The platform supports nested biomedical NER annotation and team-based review, adjudication, and final arbitration.

**Figure 2.**
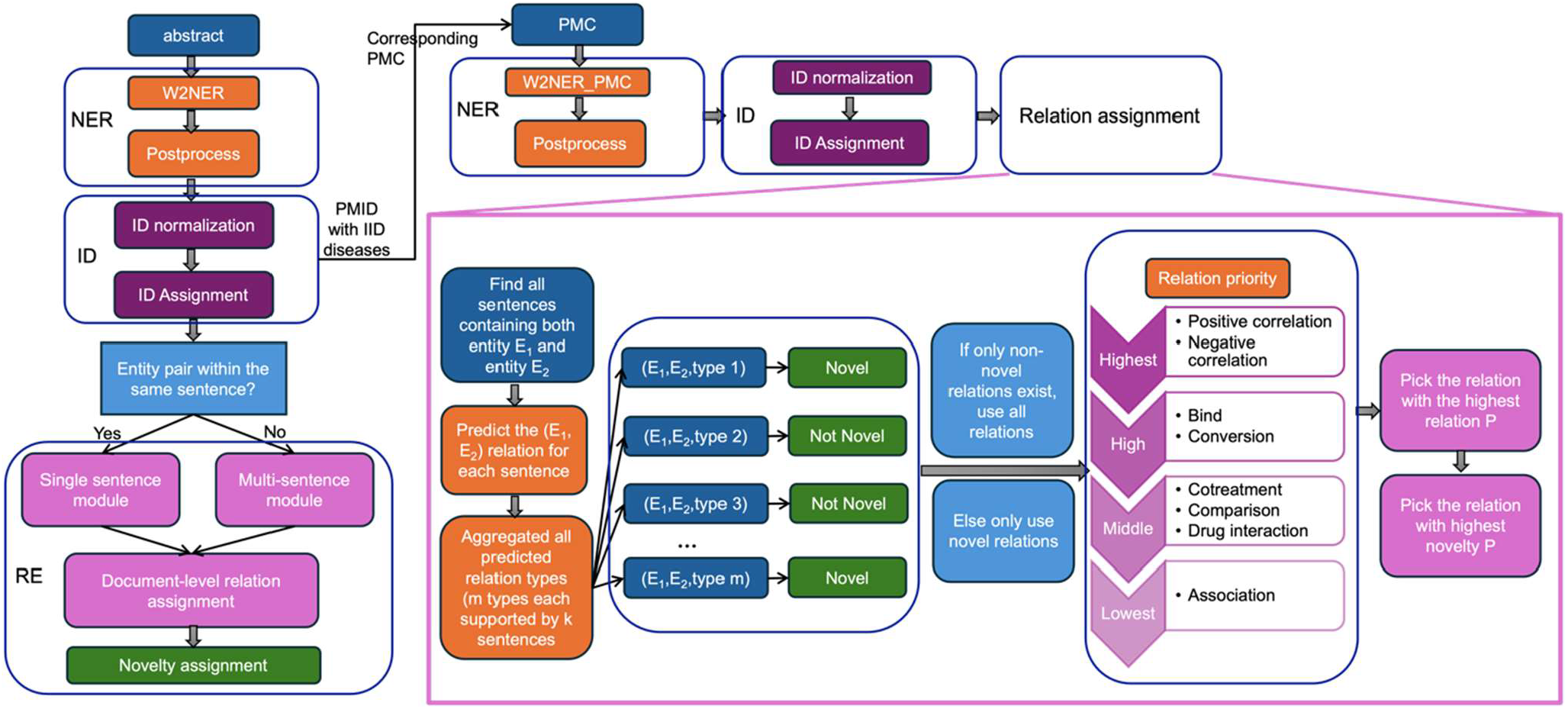
End-to-end information extraction and relation resolution pipeline for PubMed abstracts and IID-related PMC full-text articles.

**Figure 3.**
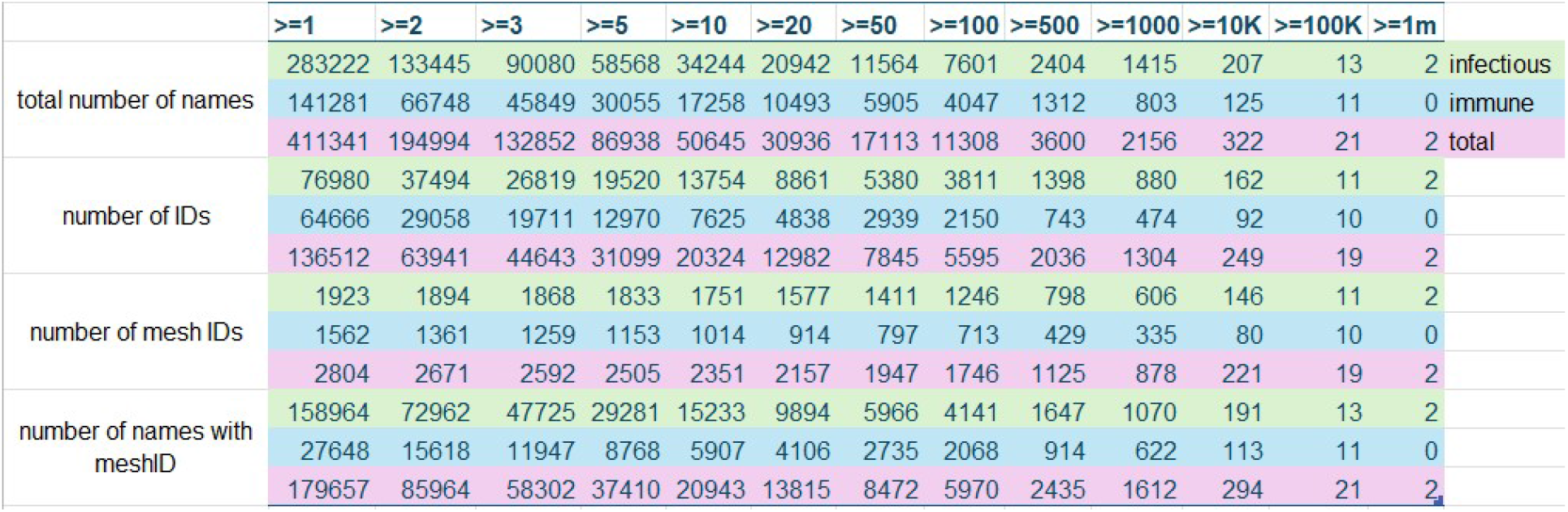
Statistical summary of ontology refinement results, shown in tabular format in the original report.

### 3.5. IID-specific knowledge graph content

Integration of large-scale entity and relation extraction outputs with the unified IID ontology produced a standalone IID-specific knowledge graph containing approximately 1,837,513 unique entities and 16,295,390 unique relations across eight relation types. The graph includes both novel and known biomedical associations with relation types covering Association, Bind, Comparison, Conversion, Cotreatment, Drug Interaction, Negative Correlation, and Positive Correlation.

The IID-specific knowledge graph was released on Zenodo under an open license. Drug repurposing code, including the PSR implementation and example workflows, was made available in the IKraph GitHub repository for application to both IKraph and the IID-specific resource.

## 4. Discussion

This work reports the construction of a dedicated knowledge graph for infectious and immune-mediated disease research by integrating large-scale literature mining, nested entity recognition, disease ontology refinement, relation extraction, and open release of the resulting resource. The key contribution is not simply an incremental update to a general biomedical KG, but the creation of an IID-focused graph whose corpus selection, disease normalization strategy, and relation integration pipeline were designed around the needs of infectious and immune-mediated disease research. The resulting graph contains approximately 1.84 million unique entities and 16.30 million unique relations, providing a structured evidence base for direct search, literature-backed association discovery, and downstream reasoning applications.

The results support the feasibility of using a domain-focused KG construction strategy at literature scale. Processing all PubMed abstracts up to April 14, 2025 allowed the pipeline to identify more than 6.3 million IID-related abstracts, from which nearly 1.4 million corresponding PMC full-text articles were retrieved and processed. This design balances breadth and depth: abstracts provide comprehensive coverage across the biomedical literature, whereas full texts provide additional evidence that may not appear in abstracts. The full-text component is particularly important for IID biology, where mechanistic details, experimental conditions, pathogen-host interactions, immune pathway measurements, and treatment-related evidence often appear outside the abstract.

The nested NER results demonstrate both the value and the difficulty of full-text biomedical extraction. The abstract-level model achieved strong in-domain performance, but its lower cross-domain performance on PMC full texts indicates that full-text articles cannot be treated as longer abstracts. Section-wise analysis further showed that model performance varies by article section, with Methods being more challenging than Results or Conclusion. The full-text model trained with combined abstract and full-text annotations improved overall robustness and showed strong external validation performance on Europe PMC annotations, especially for disease mentions. These findings justify the inclusion of full-text-specific annotation and evaluation in future KG construction projects rather than relying exclusively on abstract-trained models.

Ontology refinement was a central determinant of KG quality. The project processed 411,341 disease terms and confirmed 179,657 MeSH mappings after LLM refinement, while also incorporating manual review of high-frequency infectious and immune-related terms. This multi-tiered strategy addressed two complementary problems: maximizing recall for disease concepts encountered in large-scale NER outputs, and maintaining precision by filtering non-disease terms and correcting misleading synonym mappings. The high F1 scores achieved during noise filtering suggest that combining ontology cross-referencing with LLM-based classification can be effective when paired with expert review and post-processing. Importantly, this ontology layer also makes the KG more interoperable with external resources such as MeSH, MONDO, Disease Ontology, and HPO.

The relation extraction and resolution strategy addresses a key challenge in full-text KG construction: the same entity pair may be mentioned in multiple sentences, sections, or contexts, and different sentences may support different relation types or novelty labels. The implemented pipeline aggregates sentence-level predictions, applies novelty-based filtering, ranks relation types by informativeness, and resolves ties using probability-based criteria. This design is conservative in that it prioritizes novel relations when present while preserving non-novel evidence when no novel relation is detected. The optional abstract-level override also reflects the practical observation that abstracts often contain curated, high-salience claims that can help resolve conflicts arising from longer and more heterogeneous full-text evidence.

The IID KG has several potential uses. First, it provides a structured literature-derived map of disease-gene, chemical-disease, chemical-gene, gene-gene, variant-gene, and other biomedical relationships relevant to IID research. Second, because the graph preserves relation types and evidence provenance, users can move beyond simple co-occurrence and retrieve interpretable relation evidence. Third, the graph can support hypothesis generation through PSR-based reasoning, including indirect disease-drug and disease-gene associations for drug repurposing and target discovery. This motivated the capability using earlier BioKG/IKraph drug repurposing and target-discovery case studies; in the present work, the released IID KG and repurposing workflows provide the infrastructure for applying similar reasoning specifically to IID questions.

Several limitations should be acknowledged. The gold-standard full-text annotation set was intentionally high quality but small, consisting of eight PMC articles, which limits the diversity of full-text structures represented during model adaptation. Relation extraction remains dependent on sentence-level or sentence-pair evidence, and relations that require broader discourse-level inference may still be missed. Entity normalization also remains challenging for newly emerging disease terms, ambiguous abbreviations, and terms that fall between disease concepts, symptoms, phenotypes, organisms, and interventions. Finally, although an automated weekly update system was implemented for IKraph, routine refresh of the IID-specific KG would require adapting and validating the same update framework for the disease-focused release.

Future work should expand the full-text annotation set, improve section-aware modeling, incorporate additional relation types such as hierarchical is_a relations where appropriate, and strengthen continuous update workflows for the released IID KG. Additional validation against downstream use cases will also be important, including prospective drug repurposing, target prioritization, emerging infectious disease surveillance, and immune-mediated disease mechanism discovery. Because the KG is openly released, external users can evaluate, extend, and integrate the resource with complementary datasets, which should help refine both the graph content and its applications over time.

## 5. Conclusion

A validated, ontology-aligned IID-specific biomedical knowledge graph was constructed from large-scale PubMed abstract and PMC full-text processing. The workflow integrated gold-standard nested entity annotation, abstract and full-text NER models, a unified IID ontology, large-scale relation extraction, and relation-resolution methods. The resulting open IID KG provides a dedicated structured evidence resource for infectious and immune-mediated disease research and downstream drug-repurposing applications.

## Data and Code Availability

The IID-specific knowledge graph is publicly released on Zenodo at https://zenodo.org/records/17047896. The disease-drug repurposing code, including the PSR implementation and example workflows, is available in the IKraph GitHub repository at https://github.com/myinsilicom/IKraph/tree/main/repurposing.

## Funding and Contract Information

This work was supported by the National Institute of Allergy and Infectious Diseases of the National Institutes of Health under contract number 75N93024C00034. The content is solely the responsibility of the authors and does not necessarily represent the official views of NIAID, NIH, or HHS.

## Acknowledgments

The authors acknowledge the biomedical annotators and reviewers who contributed to gold-standard annotation, adjudication, ontology review, and pipeline validation.

